# Cytoplasmic and nuclear protein interaction networks of the synapto-nuclear messenger CRTC1 in neurons reveal cooperative chromatin binding between CREB1 and CRTC1, MEF2C and RFX3

**DOI:** 10.1101/2025.07.02.662820

**Authors:** Sylvia Neumann, Jennifer M. Achiro, Marika Watanabe, Shivan L. Bonnano, Weixian Deng, James A. Wohlschlegel, Kelsey C. Martin

## Abstract

Glutamatergic stimulation of excitatory neurons triggers the synapto-nuclear translocation of the cAMP response element (CRE) binding protein (CREB) regulated transcription coactivator 1 (CRTC1), resulting in the transcription of CREB1 target genes. Whether and how CRTC1 and CREB1 interact with other transcription factors to regulate activity-dependent transcription, and what the role of CRTC1 is in neurons beyond the activation of CREB1 regulated transcription, remains unknown. To address these questions in an unbiased manner, we used proximity labeling to identify CRTC1-proximal proteins in cytoplasmic and nuclear compartments of rodent forebrain neurons. The cytoplasmic CRTC1 proxisome included a variety of signaling pathways and downstream cellular processes involved in synaptic plasticity. In contrast, the nuclear CRTC1 proxisome included transcription factors that mediate activity-dependent transcription, chromatin factors, and splicing factors. Our data revealed that CRTC1 and CREB1 interact with MEF2C and RFX3 transcription factors in an activity-dependent manner. Thus, in chromatin immunoprecipitation-sequencing experiments, CREB1 was prebound to chromatin regions containing bZIP motifs in a manner that was unchanged by neuronal activity, while glutamatergic stimulation triggered the recruitment of CRTC1 and CREB1 to activity-dependent enhancers enriched in motifs for MEF2C and RFX3. Collectively, these results not only enhance our understanding of the role of cytoplasmic and nuclear CRTC1 in neurons, but also reveal a role for CRTC1 in promoting cooperativity of CREB1 with other transcription factors in response to synaptic activity.

## Introduction

Experience-dependent changes in synaptic efficacy underlying learning and memory require changes in gene expression for their persistence^1–3^. The highly polarized and compartmentalized morphology of neurons presents a challenge to the regulation of gene expression, raising the question of how synaptic signaling and activity-dependent changes in gene expression are spatially coordinated. Neuronal stimulation activates several synapto-nuclear signaling pathways, including electrochemical signaling, calcium wave propagation from distal sites to the nucleus, and regulated active nuclear protein import^4^. One prominent synapto-nuclear messenger that undergoes activity-dependent nuclear import is the cAMP response element (CRE) binding protein (CREB) regulated transcription coactivator 1 (CRTC1)^5,6^. Indeed, in mature neuronal cultures, CRTC1 was the most enriched protein that translocated to the nucleus following glutamatergic stimulation^7^.

In mammals, CRTCs are expressed as three different isoforms, with CRTC1 being highly expressed in neurons. In silenced neurons, CRTC1 is highly phosphorylated^5,6^ and is anchored at synapses by binding to synaptically-localized 14-3-3ε proteins^6^. Following synaptic stimulation, an increase in intracellular Ca^2+^ leads to the dephosphorylation of CRTC1 by calcineurin and its release from 14-3-3ε^6^. This allows for nuclear import of CRTC1, binding to the basic leucine zipper (bZIP) domain of CREB1^8,9^ and the cooperative recruitment of general transcription and chromatin factors, such as Taf_II_130^8^, CBP and BRD4^10–12^, resulting in the transcription of CREB1 target genes^12–16^. CRTC1 also contains a nuclear export signal, and nuclear export and rephosphorylation of CRTC1 by SIK triggers the export of CRTC1 to the cytoplasm^17,18^.

Many lines of evidence have demonstrated a role for CREB1 in the conversion of short-term to long-term plasticity and memory^19–21^. Likewise, CRTC1 has been shown to regulate the expression of genes important for synaptic plasticity and to play a role in long-term potentiation and memory formation^6,13–16,22–26^. Moreover, dysregulation of CRTC1/CREB1 mediated transcription in neurons has been implicated in neurodegenerative diseases such as Alzheimer’s, Parkinson’s and Huntington’s as well as in psychiatric disorders^27,28^.

Little is known about the function of CRTC1 in neurons beyond co-activation of CREB1-regulated transcription. However, in other cell types, CRTC1 has been shown to coactivate transcription in coordination with other factors. In HeLa cells, CRTC1 was shown to activate transcription regulated by the bZIP transcription factor AP-1^29^, which consists of a dimer of cFOS and cJUN family members^30^. Similarly, genetic interactions demonstrated a role for CRTC1 in AP-1 regulated transcription in *C. elegans* neurons. The same study also demonstrated an interaction of CRTC1 with other chromatin factors, including the histone methyltransferase SET1 and WD40 domain-containing proteins, which are important for regulation of life span^31^. The CRTC family member CRTC2 has been shown to bind the bZIP transcription factor ATF6 to regulate the expression of ATF6 target genes under conditions of ER stress^32^. CRTC2 has also been implicated in another stress pathway by directly binding to and mediating the co-operative activity between CREB1 and the glucocorticoid receptor in regulating gluconeogenesis^33^. Other studies in non-neuronal cells have reported roles for CRTCs outside of transcription, including involvement in splicing and membrane trafficking^34,35^. These findings suggest that CRTC1 may have undiscovered functions in neurons.

To examine, in an unbiased manner, the function of CRTC1 during activity-dependent neuronal plasticity we utilized engineered ascorbic acid peroxidase 2 (APEX2) proximity mapping with mass spectrometry^36,37^ to identify the CRTC1 proxisome, and utilized Chromatin immunoprecipitation (ChIP)-sequencing (ChIP-seq) to generate genome-wide CRTC1 and CREB1 chromatin binding maps in silenced and stimulated neurons. We found that cytoplasmically-localized CRTC1 interacted with key pathways supporting synaptic function, such as membrane trafficking, translation, and small GTPase signaling, whereas nuclear-localized CRTC1 was part of a network of chromatin and transcription factors, including MEF2 and RFX transcription factors, as well as a network of proteins regulating alternative splicing and posttranscriptional mRNA processing. We demonstrate that CRTC1 and CREB1 bind regions of active chromatin in an activity-dependent manner, especially in regions with MEF2 and RFX motifs. Furthermore, we used proximity ligation assays to visualize the interaction of CRTC1 and CREB1 proteins with MEF2C and RFX3. Together, our findings indicate that CREB1 coordinates with CRTC1, MEF2C and RFX3 to regulate transcription in an activity-dependent manner in neurons.

## Results

### Identification of CRTC1 proximal proteins in silenced and stimulated neurons

To investigate the cellular pathways associated with CRTC1’s role in activity-dependent neuronal function, we utilized a proximity labeling strategy to identify the cytoplasmic and nuclear CRTC1 proxisome. In cultured rat forebrain neurons, we expressed CRTC1 fused to APEX2-FLAG under the control of the neuron-specific synapsin promoter. We transduced forebrain neurons with lentiviral particles encoding the CRTC1-APEX2 fusion protein at 15 days in vitro (DIV). To investigate the proxisome of cytoplasmic versus nuclear localized CRTC1, we either silenced neurons at DIV 22 with 1 µM tetrodotoxin (TTX) for 1 hour or stimulated neurons with 40 µM bicuculline (BIC) for 3 hours while inhibiting nuclear export with Leptomycin B (LMB)^38–40^. During the last 30 minutes of treatment, cultures were incubated with biotin-phenol followed by a 1 min pulse of H_2_O_2_ to induce APEX2-mediated biotinylation of CRTC1 proximal proteins prior to processing for mass spectrometry analysis (Figure 1A). As controls, untransduced neurons were processed identically.

**Figure 1:**
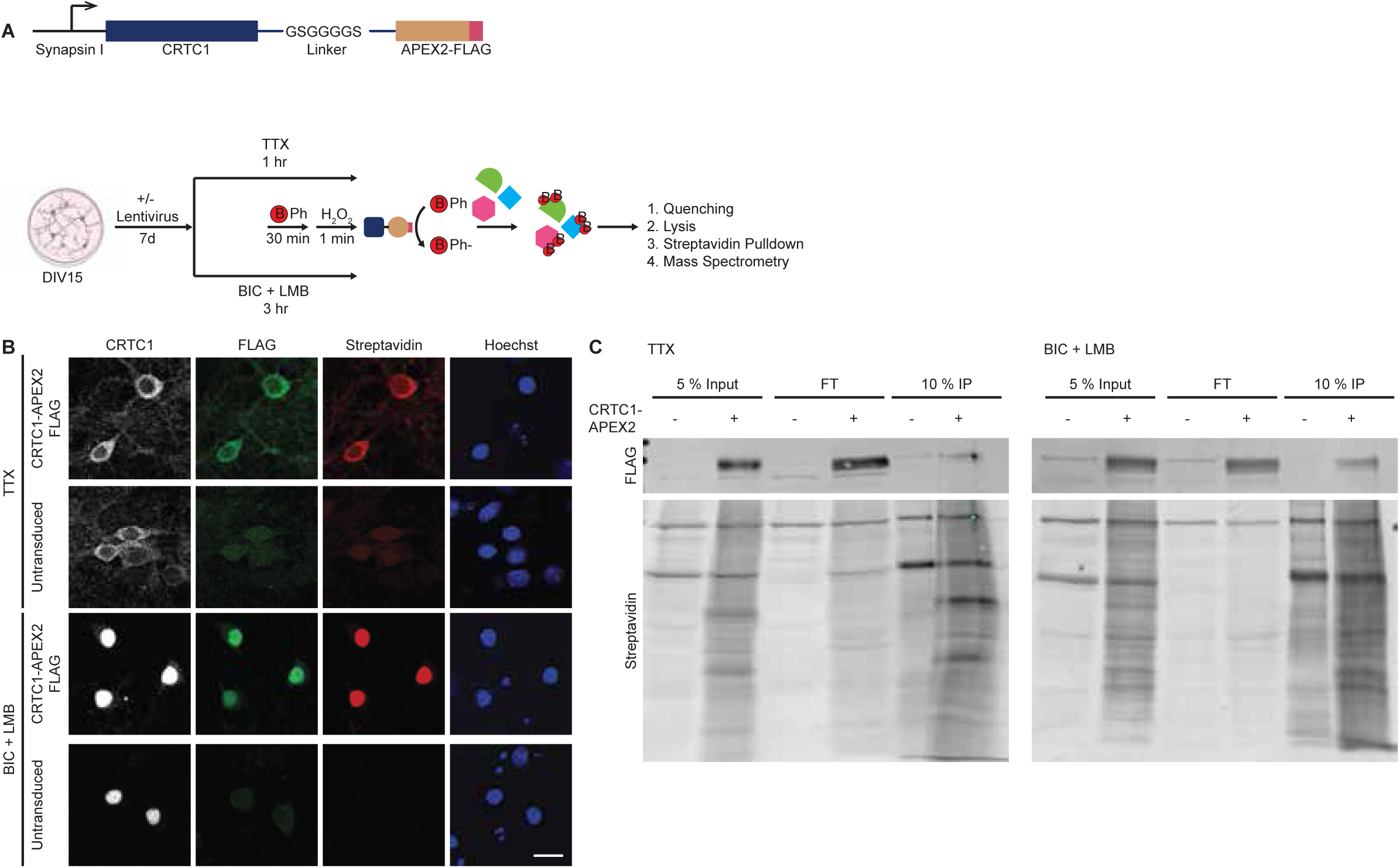
Proximity ligation labeling of cytoplasmic and nuclear CRTC1: A) Experimental Design: Upper panel: Schematic of pLSyn-CRTC1-APEX2-FLAG construct. Lower panel: Schematic of the experimental design (created in Biorender: https://Biorender.com/zqfz6qp.) B) Fluorescent images of silenced (TTX) or stimulated (BIC + LMB) neurons either untransduced or expressing CRTC1- APEX2, stained for CRTC1, FLAG, biotinylated proteins using streptavidin or DNA using Hoechst dye. C) Western Blot analysis of biotinylated proteins in cell lysates (input), flow-through (FT) or streptavidin precipitated eluates (IP) in silenced (TTX) or stimulated (BIC + LMB) neurons either untransduced or expressing CRTC1-APEX2, as detected by anti- FLAG antibodies or streptavidin staining. Scale bar represents 10 µm.

Similar to endogenous CRTC1^6^, CRTC1-APEX2 localized to the cytoplasm in neurons silenced with TTX, and CRTC1-proximal proteins detected by fluorescently labeled streptavidin were restricted to the cytoplasmic compartment (Figure 1B). Conversely, neuronal stimulation with BIC + LMB resulted in robust nuclear accumulation of CRTC1-APEX2, and biotinylated proteins (detected by streptavidin labeling) localized to the nuclear compartment (Figure 1B). Control untransduced neurons were negative for FLAG and streptavidin labeling (Figure 1B). Western blot analysis detected proteins biotinylated by CRTC1-APEX2 in the TTX and BIC + LMB conditions in transduced neurons both in the input and in streptavidin immunoprecipitation (IP) samples, whereas biotinylated proteins in untransduced neurons were limited to those known to be endogenously biotinylated^36^ (Figure 1C). IP-isolated proteins from transduced and untransduced neurons from both TTX and BIC + LMB conditions were processed for mass spectrometry and quantified using label free methods.

### Proximity-labeled proteins indicate a role of CRTC1 in various cellular pathways important for synaptic function

In silenced neurons, we identified 1550 proteins enriched in transduced compared to untransduced neurons (Figure 2A). We first determined the subcellular localization of cytoplasmic CRTC1 proximal proteins using the human annotation of the COMPARTMENTS database^41^. Consistent with a cytoplasmic localization of CRTC1 in silenced neurons, ∼98% of identified proximal proteins were classified as cytoplasmic proteins or proteins that localize both in the cytoplasm and the nucleus (Figure 2B).

**Figure 2:**
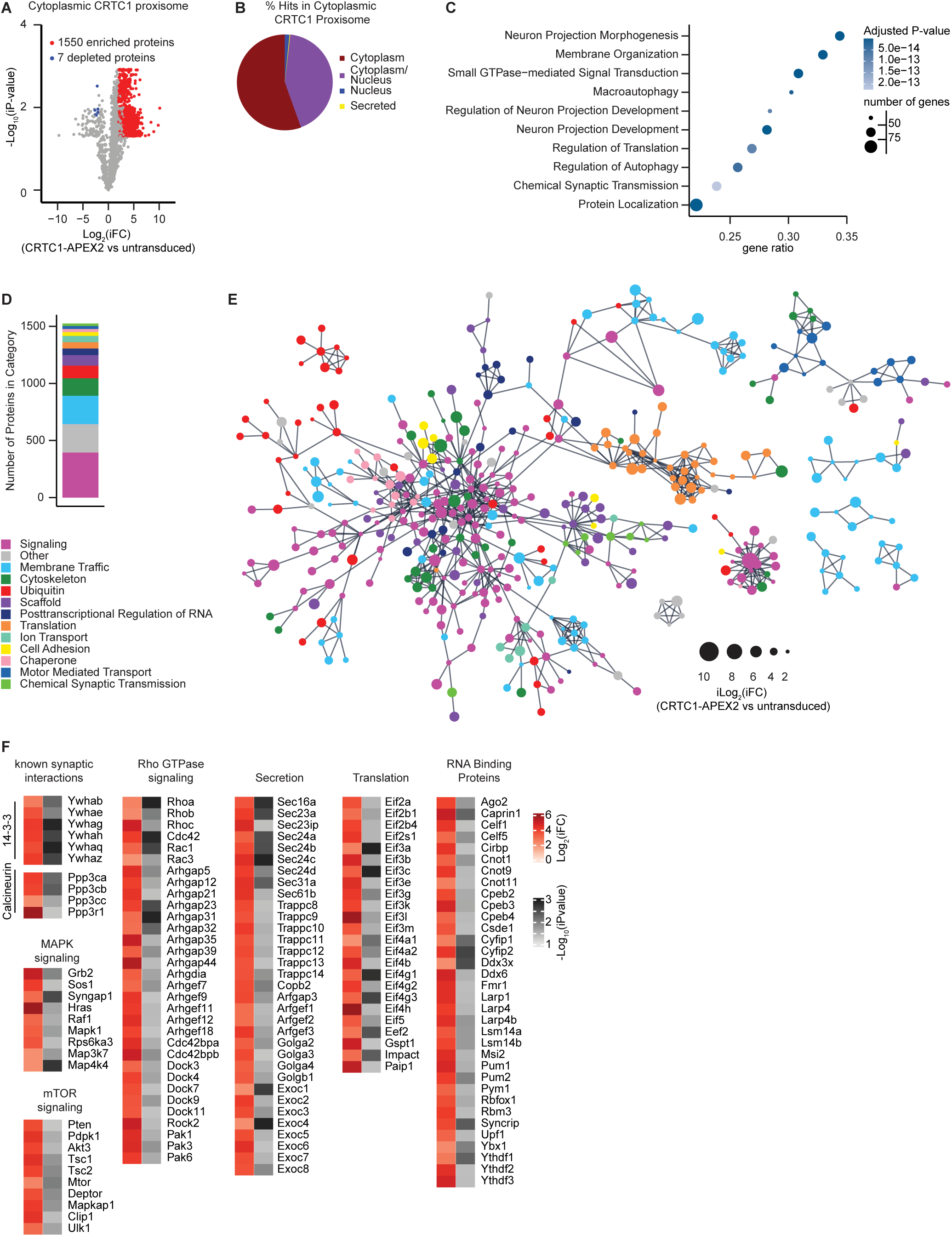
CRTC1 proxisome in silenced neurons: A) Volcano plot of cytoplasmic CRTC1 proximal proteins. Proteins enriched in CRTC1-APEX2 over untransduced (Log_2_ iFold Change > 2 and iP-Value <0.05) are depicted in red, depleted proteins in blue and proteins that were not enriched in grey. B) Pie chart of subcellular localization of CRTC1 proximal proteins detected in silenced neurons. Subcellular localization was obtained from COMPARTMENTS Annotation and simplified into shown categories. C) Gene ontology enrichment analysis for Biological Process of cytoplasmic CRTC1 proximal proteins dobtained from EnrichR, with the size of each marker indicating the number of genes in proxisome for that category and shading indicating adjusted p-value. D) Number of cytoplasmic CRTC1-proximal proteins in functional categories obtained from STRING database. E) Known protein interactions of cytoplasmic CRTC1 proximal proteins obtained from the STRING database, with marker size for each protein indicating enrichment in CRTC1 proxisome. F) Heatmaps of CRTC1 proxisomal proteins showing Log_2_ of imputed Fold Change and negative Log_10_ of imputed P-values.

Gene ontology analysis using Enrichr^42–44^ revealed cytoplasmic CRTC1 proximal proteins were enriched in pathways important for synaptic function, such as synaptic chemical transmission and synapse organization, neuron projection morphogenesis and development, translation, and signaling pathways, in particular small GTPase signaling (Figure 2C). To further functionally characterize cytoplasmic CRTC1 proximal proteins, we manually annotated protein function guided by Genecards^45,46^ and STRING descriptions^47^ (Figure 2D) and visualized known high confidence interactions between these proteins using STRING and Cytoscape^48^ (Figure 2E). Annotation and visualization of the generated interactome revealed a large cluster of proteins involved in cellular signaling as well as proteins involved in vesicular trafficking and in translation.

As expected, APEX2 labeling identified members of the 14-3-3 family of scaffolding proteins and the subunits of the phosphatase calcineurin, PPP3CA-C and its regulatory subunit PPP3R1, which are known to regulate CRTC1 in neurons^6^ (Figure 2F). Detailed examination revealed association of CRTC1 with several interconnected signaling networks, including prominent components of the Ras/MAPK, mTOR and Rho-GTPase signaling cascades. Signaling by these networks is essential for synaptic function and has been shown to be upstream of CRTCs^35,49^. We also found upstream and downstream components of the aformentioned signaling networks as shown in Figure 2F.

The aforementioned signaling cascades signal to various cellular processes that support structural changes at the synapse, such as membrane trafficking, translation of new soluble and membrane proteins, and signaling to the nucleus to regulate transcription. Ubiquitously expressed CRTC2 has been shown to be phosphorylated by mTOR directly regulating ER-to-Golgi trafficking in hepatocytes^35^. Similarly, we identified CRTC1-proximal structural and regulatory proteins that function along the secretory pathway, such as SEC23, SEC24 and SEC31, TRAPPC, GOLGA and EXOC subunits (Figure 2F). MAPK and mTOR also signal to translation initiation and elongation factors to regulate local activity-dependent translation of synaptic proteins^50^. Notably, we identified a considerable number of translation initiation factors and RNA binding proteins (or their interaction partners) that regulate many aspects of translation (Figure 2F). Importantly, most of these proteins, such as CPEBs, CYFIPs, FMRP, PUMs and RBFOX1, are key regulators of neuronal transcripts, and mutations in these proteins lead to neurodevelopmental and neuropsychiatric disorders^51–55^.

Collectively, we found that the cytoplasmic CRTC1 proxisome includes proteins that are known synaptic interaction partners of CRTC1 as well as novel CRTC1 interactions. The identification of cytoplasmic CRTC1-proximal signaling, membrane trafficking and translation-related proteins in silenced neurons reveals possible unexplored synaptic functions of CRTC1 in shaping synaptic plasticity.

### Proximity labeling places CRTC1 in a larger network of transcription factors, transcriptional regulators, and splicing factors

In stimulated neurons, we identified 580 proximal proteins enriched in CRTC1-APEX2 expressing neurons compared to untransduced neurons (Figure 3A). Analysis of the subcellular localization of CRTC1 proximal proteins in stimulated neurons confirmed that ∼ 93% of identified proximal proteins localized either to the nucleus or both the nucleus and the cytoplasm (Figure 3B), in agreement with the activity-dependent nuclear localization of CRTC1. Gene ontology analysis revealed that nuclear CRTC1 proximal proteins were enriched in biological processes related to transcription, RNA processing and splicing (Figure 3C). Similarly, manual annotation and high confidence protein interactions of nuclear CRTC1 proximal proteins revealed interactome clusters related to transcription, splicing and RNA processing, and also highlighted proteins involved in epigenetic regulation and ubiquitin related pathways (Figure 3D,E).

**Figure 3:**
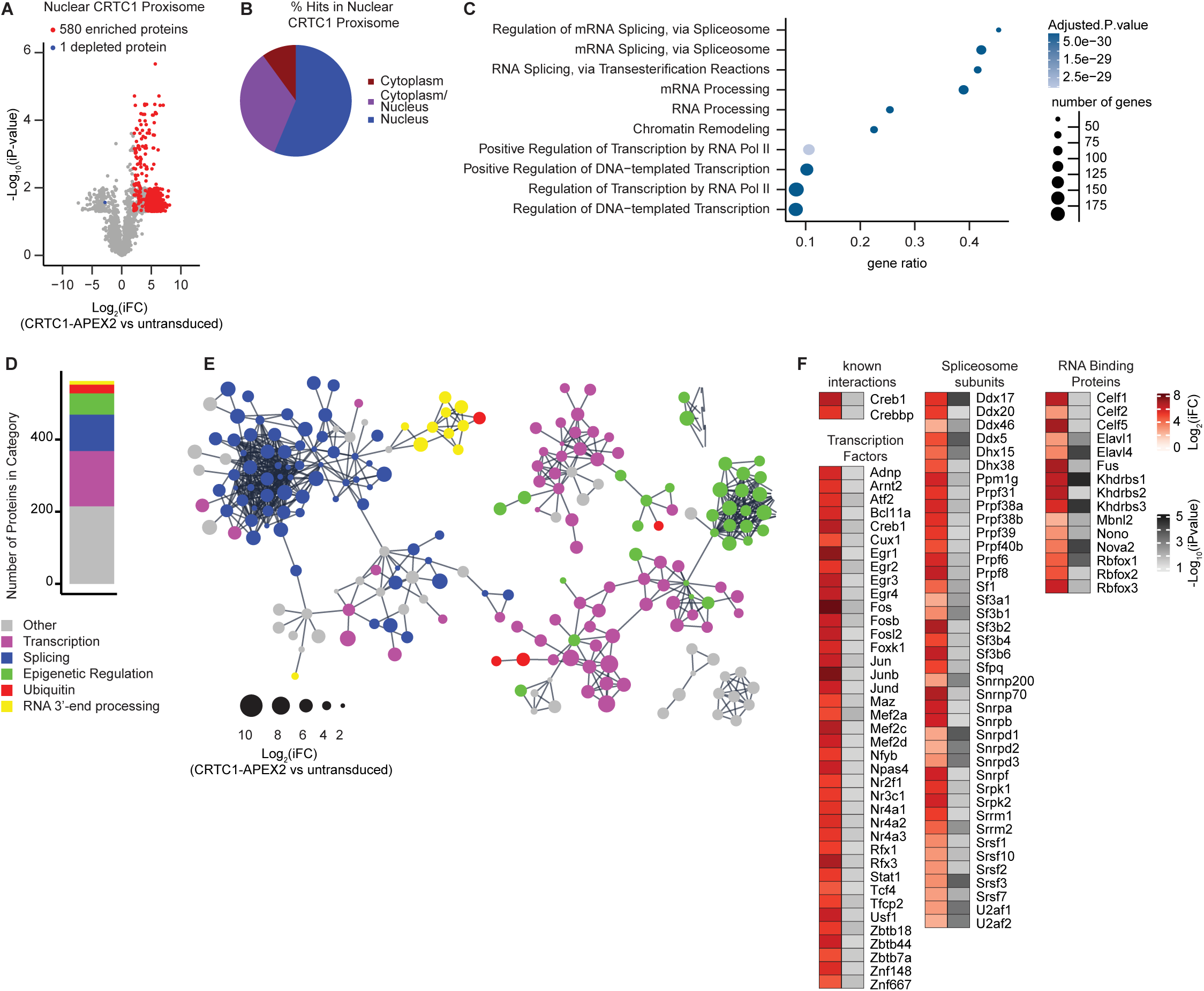
CRTC1 proxisome in stimulated neurons: A) Volcano plots of nuclear CRTC1 proximal proteins. Proteins enriched in CRTC1-APEX2 over untransduced (Log_2_ iFold Change > 2 and iP-Value <0.05) are depicted in red, depleted proteins in blue and proteins that were not enriched in grey. B) Pie chart of subcellular localization of CRTC1 proximal proteins detected in stimulated neurons. Subcellular localization was obtained from COMPARTMENTS Annotation and simplified into shown categories. C) Gene ontology enrichment analysis for Biological Process of CRTC1 proximal proteins detected in stimulated neurons obtained from EnrichR. D) Number of proteins in functional categories obtained from STRING database. E) Known protein interactions of CRTC1 proximal proteins detected in stimulated neurons obtained from the STRING database. F) Heatmaps of CRTC1 proxisomal proteins showing Log_2_ of imputed Fold Change and negative Log_10_ of imputed P-values.

In the nucleus, CRTC1 has been reported to interact with CREB1^8,56^ and CBP^10,11^, which we also identified as nuclear CRTC1-proximal proteins in our experiments (Figure 3F). In addition to CREB1, we identified several other transcription factors known to be involved in activity-dependent transcription. These include other bZIP containing factors, such as ATF2, and members of the FOS and JUN families that form the transcription factor AP-1, known to interact with CREB1 and CRTC1^29^. We also identified other factors from different transcription factor families, namely members of the EGR, NR4A, MEF2 and RFX families, as well as ARNT2 and NPAS4, which belong to the bHLH-PAS family (Figure 3F). Like CREB1, MEF2 transcription factors are regulated by synaptic activity mainly at the level of posttranslational modifications, such as phosphorylation, acetylation, sumoylation, cleavage and S-nitrosylation^57^. MEF2C and CREB1 have been shown to cooperate at activity-dependent enhancers in neurons stimulated with the neuromodulator Reelin^58^. RFX factors are important for ciliogenesis and have recently been implicated in neurodevelopmental disorders^59^. Moreover RFX3, which we identified as a CRTC1-proximal protein, has been implicated in the regulation of CREB1 mediated transcription^60–62^.

A second major group of proteins identified as proximal to nuclear CRTC1 were involved in RNA binding and splicing. These include core components of the spliceosome (Figure 3F) and RNA binding proteins (RBPs) that are known to regulate various aspects of mRNA processing in neurons, including alternative splicing^63–65^. Of these, we identified members of the CELF, ELAVL, KHDBRBS and RBFOX families (Figure 3F). Further, we found NOVA2 and MBNL2 in the CRTC1 proxisome, RBPs that have important functions in the nervous system^63^. We also identified NONO (p54NRB), which has previously been shown to bind to CRTC1 in non-neuronal cells in a manner that regulates transcription^66^. NONO also functions in splicing, and CRTC1 was also reported to play a role in alternative splice site selection of CREB1 target genes^34^. Notably, some CRTC1-proxisomal RBPs also regulate mRNAs in the cytoplasm^64,65^. Interestingly, we identifed the RBPs RBFOX1, CELF1 and CELF5 proximal to CRTC1 in TTX-silenced neurons (Figure 2F), indicating that they might function with CRTC1 to regulate posttranscriptional aspects of mRNA biology. Together, these data suggest that, in addition to binding CREB1, neuronal CRTC1 functions in the nucleus cooperatively with activity-dependent transcription factors such as ATF, FOS, JUN, EGR, NR4A, MEF2 and RFX. Furthermore, these data point to a novel role for CRTC1 in the activity-dependent regulation of splicing in neurons.

### CRTC1 and CREB1 bind to chromatin regions that are enriched in MEF2C and RFX3 motifs

To further investigate whether CRTC1 and CREB1 cooperate with other transcription factors to regulate activity-dependent gene expression, we performed ChIP-seq for CRTC1 and CREB1 in neurons along with ChIP-seq for the histone mark histone 3 lysine 4 trimethylation (H3K4me3), an epigenetic marker that is found primarily in regions proximal to transcription start sites of actively transcribed genes^67,68^, and ChIP-seq for histone 3 lysine 27 acetylation (H3K27ac), an epigenetic marker that is associated with the activation of transcription and found in both enhancer and promoter regions of actively transcribed genes^67,69^. Neurons were either stimulated with 40 µM BIC for 15 min or silenced with 1 µM TTX for 30 min before fixation. Unlike CREB1, CRTC1 is not known to bind directly to chromatin, and we therefore utilized disuccinimidyl glutarate (DSG) in addition to formaldehyde to crosslink proteins to chromatin before IP^70^. Summits of ChIP-seq peaks from four biological replicates per condition were determined using model-based analysis for ChIP-seq (MACS2)^71^. To focus on genomic regions that are associated with active transcription, we filtered CREB1 bound-regions to include only those also displaying H3K27ac. Using these criteria, we identified 34,185 consensus regions for CREB1 in BIC that we further analyzed for CRTC1 binding, genomic annotation, motif occurrence and activity-dependent binding.

Analysis of CREB1 and CRTC1 ChIP-seq binding profiles showed that CREB1 was bound to chromatin in both silenced and stimulated neurons, as previously reported^72–74^ (Figure 4A, left panel). In contrast, CRTC1 was recruited to regions of CREB1 binding only in stimulated neurons, consistent with its activity-dependent nuclear translocation (Figure 4A, right panel). We found that most CREB1-bound regions were annotated to intergenic and intronic regions with about ∼20% of CREB1-bound regions annotated to promoters (Figure 4B). Figure 4C shows CREB1 and CRTC1 binding at the locus of the activity-dependent gene *cFos,* illustrating the activity-dependent binding of CRTC1 to the sites of prebound CREB1 at the promoter and upstream enhancer regions. In the stimulated condition (BIC), we observed the CRTC1/CREB1 complex at promoters, marked by the presence of H3K4me3 and H3K27ac, and at distal regions, such as intronic and intergenic regions, marked by H3K27ac (Figure 4D). These results indicate that CRTC1 is recruited to regions of chromatin with CREB1 binding in proximal promoter and distal enhancer regions after synaptic activity.

**Figure 4:**
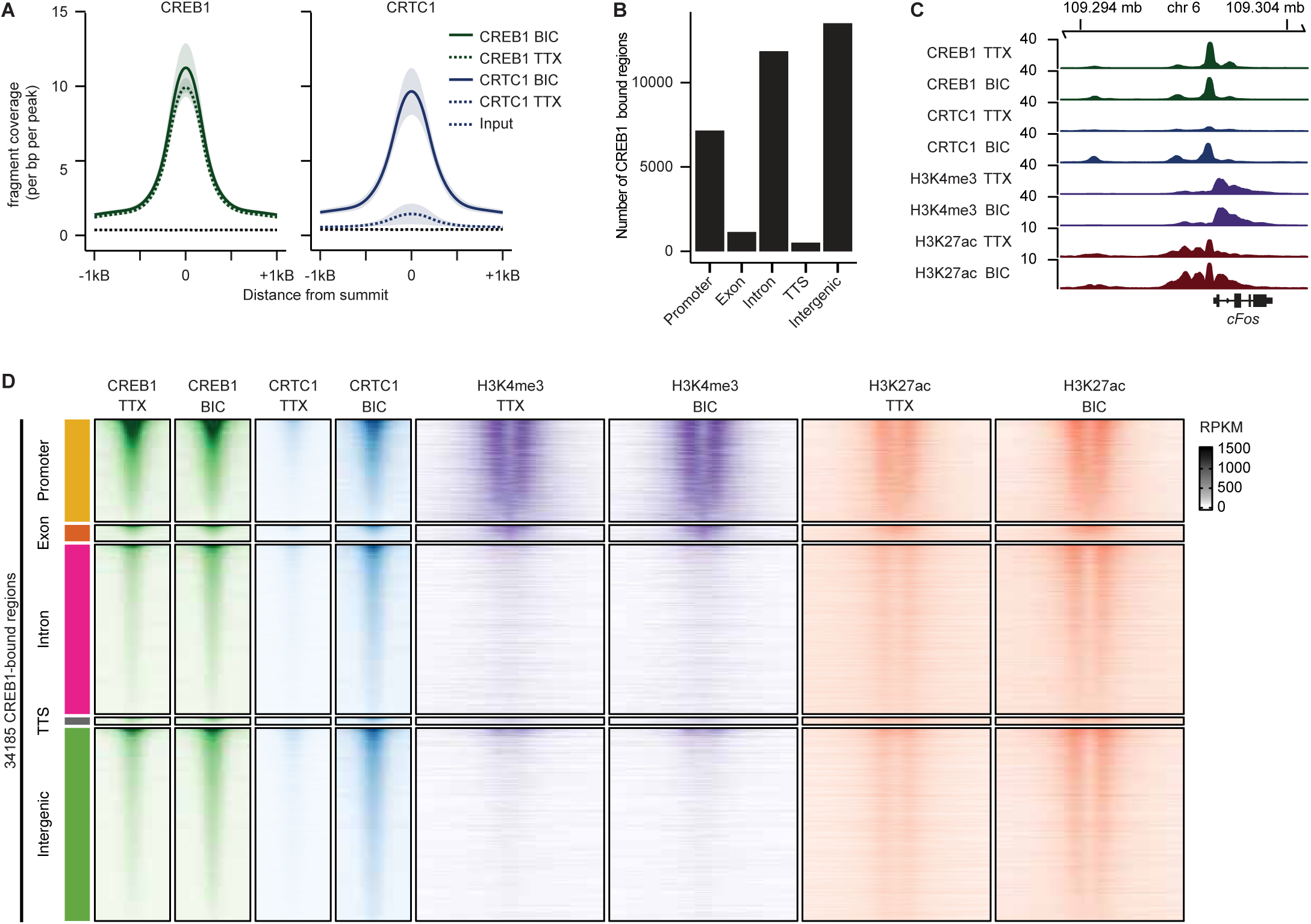
Activity-dependent binding of CRTC1 to regions of CREB1 binding at proximal and distant genomic regions. A) ChIP-seq profiles of CREB1 and CRTC1 in regions of CREB1 binding in stimulated neurons (34,185 regions). Solid and dashed lines represent average normalized ChIP-seq signal of four replicates for each condition and shading represents standard deviation. B) Genomic annotation of CREB1 consensus binding regions in stimulated neuronal cultures. C) Browser tracks of CREB1, CRTC1, H3K4me3 and H3K27ac ChIP-seq signal in stimulated and silenced neuronal cultures. Panels show the average normalized ChIP-seq signal at the c*Fos* locus. D) Heatmaps of average normalized ChIP-seq signals for CREB1, CRTC1, H3K4me3 and H3K27ac in stimulated (BIC) and silenced (TTX) neurons. Normalized ChIP-seq signals are plotted between +/-1 kB from the peak center for CREB1 and CRTC1 or +/- 2.5 kB for H3K4me3 and H3K27ac flanking the center of CREB1 binding regions in stimulated neuronal cultures. RPKM = reads per kilobase per million mapped reads.

Motif analysis of CREB1-bound regions (MOtif aNAlysis using Lisa: monaLisa)^75^ revealed 409 statistically significant motifs (adjusted p-value < 0.05), the most significant of which belonged to the bZIP factor proteins CREB1, ATF1, AP1, FOS and JUN (Figure 5A). Furthermore, we found that motifs for transcription factors of the MEF2 family and the RFX family such as MEF2A/C and RFX1/2/3/4/5 were also enriched in CREB1-bound regions. Comparing motif enrichment across different genomic regions revealed that motifs for CREB1 and other bZIP factors were significantly enriched in regions annotated to promoters and to a lesser extent to introns (Figure 5B). In contrast, motifs for MEF2 and RFX transcription factors were depleted in promoters but enriched in intronic and intergenic regions. We found that CREB1 motifs co-occurred with either MEF2C or RFX3 motifs in about a fourth of CREB1-bound regions, however it was rare for the motifs for all three transcription factors to be present in the same region (Figure 5C). Together with our findings that bZIP, MEF2 and RFX proteins were present in the nuclear CRTC1 proxisome, the enrichment of these proteins’ motifs in CREB1-bound chromatin regions indicate that the CRTC1/CREB1 complex may coordinate with these transcription factors during activity-dependent transcription in neurons. Furthermore, these results suggest that CREB1 co-binds with bZIP transcription factors in promoters, but with MEF2 or RFX proteins at enhancers.

**Figure 5:**
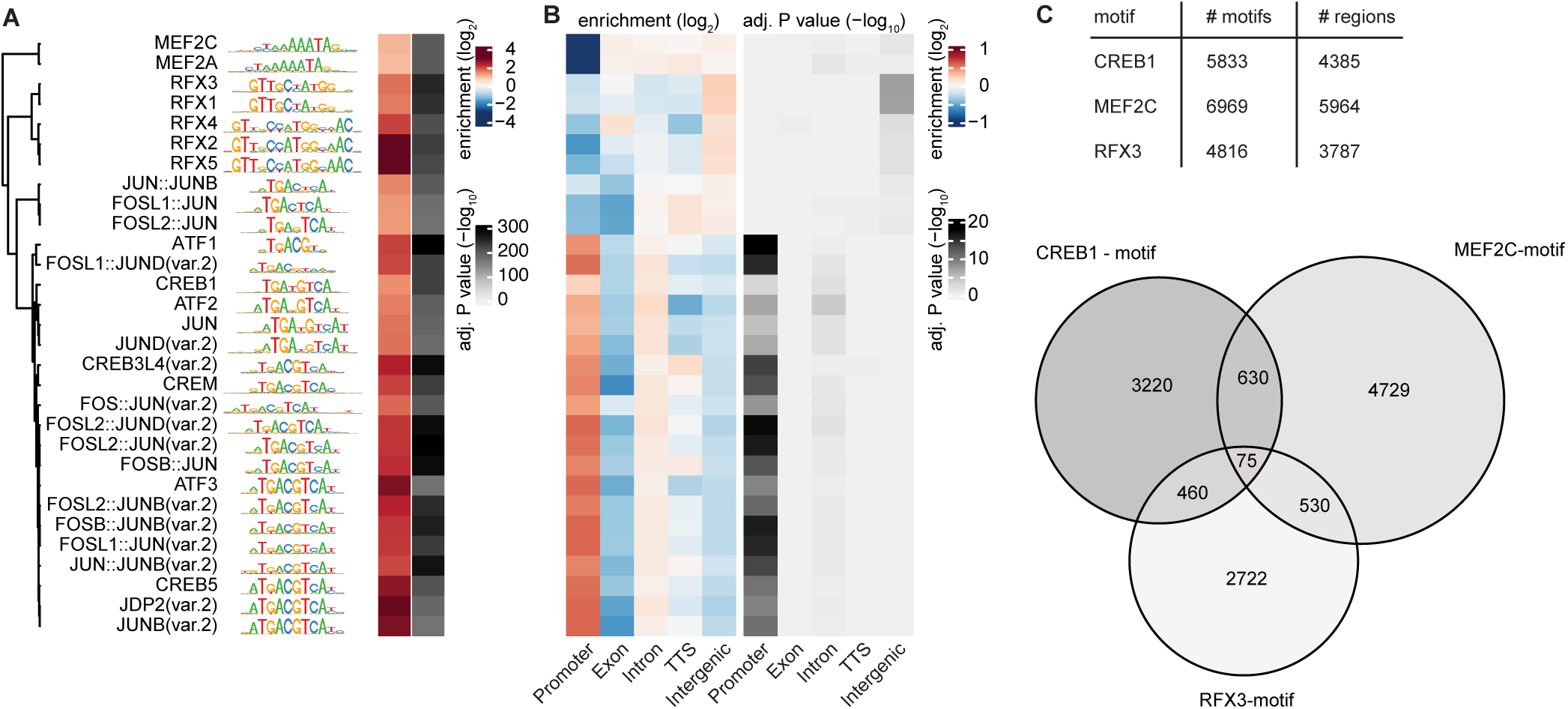
CRTC1/CREB1 binding regions are enriched for bZIP factor, MEF2C and RFX3 motifs: A) MOtif aNAlysis with Lisa (monaLisa) showing the top 30 most significant motifs identified in CREB1 binding regions. B) Motif enrichment in peak sets classified by genomic annotation. C) CREB1 binding regions were scanned for instances of motifs. Table: number of motif occurrences and corresponding number of regions. Lower panel: Overlap of CREB1 binding regions containing motifs for CREB1, MEF2C and RFX3.

To further validate our findings, we performed ChIP-seq for CRTC1 in the CA1 region of mouse hippocampus. For these experiments, acute hippocampal slices were prepared from adult mice and treated with either artificial cerebrospinal fluid (ACSF) as a control or with 100 µM dihydroxylphenolglycine (DHPG) for 10 min, which leads to robust nuclear translocation of CRTC1 in hippocampal CA1 (Figure 6A) and induces long-term depression^76–79^. Similar to our observations in cultured neurons, we found that CRTC1 was recruited to promoter and enhancer regions at the *cFos* locus in CA1 of stimulated slices (Figure 6B) and primarily bound to promoters, intronic and intergenic regions (Figure 6C, D). Corroborating our findings in cultured neurons, motif analysis for CREB1, MEF2C and RFX3 motifs revealed enrichment for these motifs in regions of CRTC1 binding in hippocampal CA1 (Figure 6E).

**Figure 6:**
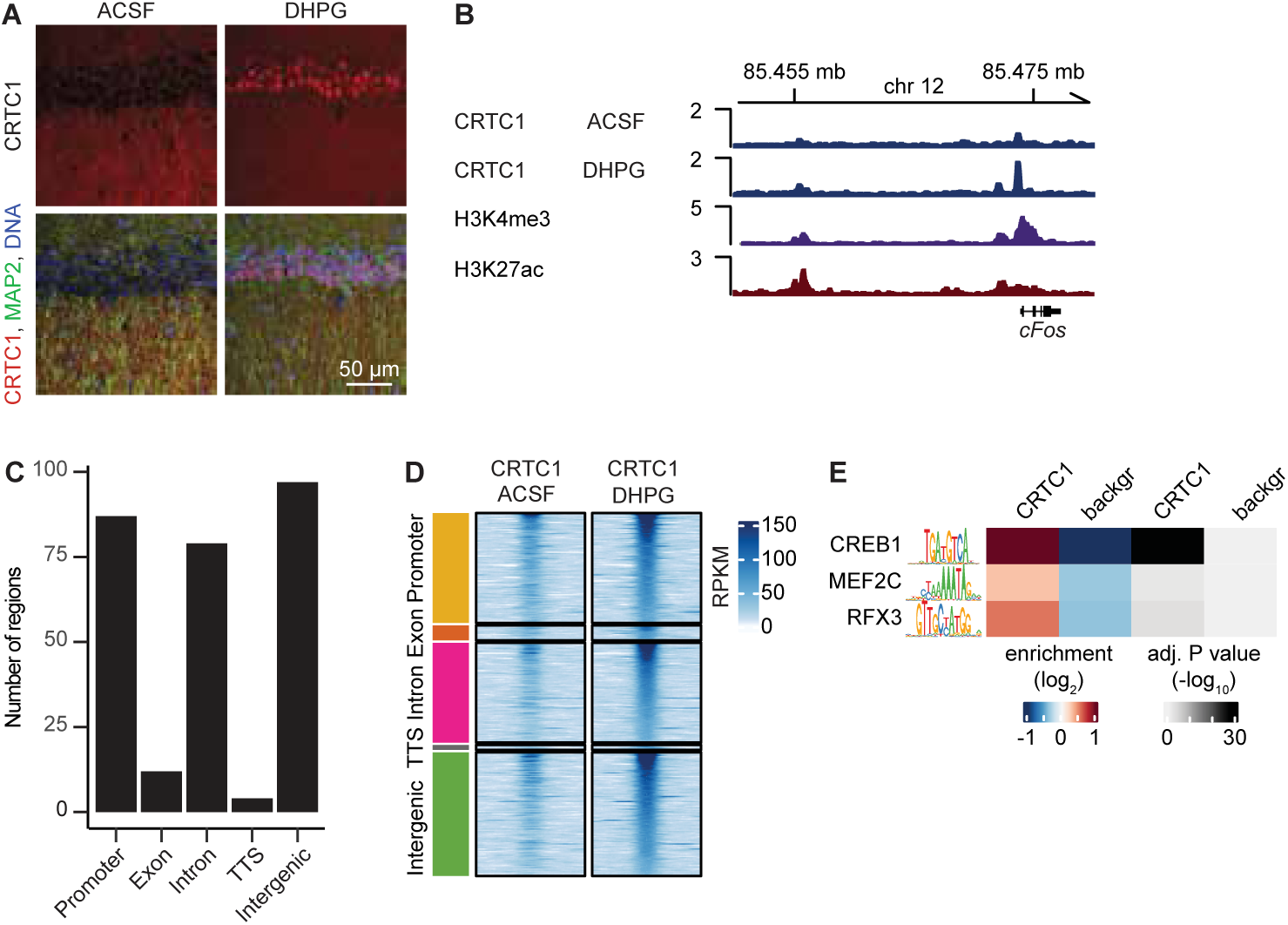
CRTC1 binds to regions enriched in motifs for MEF2C and RFX3 in hippocampus: A) Images of CRTC1 staining in the hippocampal CA1 region in control (ACSF) or stimulated (DHPG) conditions. B) Browser tracks of ChIP-seq signal from hippocampal CA1 at the *cFos* locus of CRTC1 with ACSF or DHPG. ChIP-seq signal for H3K4me3 and H3K27ac from the CA1 region of naïve animals were obtained from study GSE74971. C) Genomic annotation of CRTC1 consensus peaks from hippocampal CA1. D) Heatmaps of average normalized ChIP-seq signals for CRTC1 from CA1 treated with ACSF or DHPG. E) Motif enrichment for CREB1, MEF2C and RFX3 in CA1 CRTC1 binding regions.

### Activity-dependent chromatin binding of CREB1 occurs in regions enriched for MEF2C and RFX3 motifs

We performed differential binding affinity analysis using DiffBind^80^ and identified 10,112 regions that showed a significant increase in binding affinity of CREB1 to chromatin in BIC-stimulated neurons compared to TTX-silenced neurons (Figure 7A). We found that few regions showed decreased CREB1 binding (266 regions) with activity. Interestingly, regions that showed an increase in binding affinity had low CREB1 binding in TTX-silenced neurons but gained CREB1 binding with BIC-stimulation (Figure 7A, B, left panel). Furthermore, H3K27ac upstream and downstream of these binding events also significantly increased with stimulation, indicating that CREB1 binding affinity increased at activity-dependent enhancers (Figure 7A, B, right panel). Importantly, average CRTC1 recruitment was higher at activity-dependent regions compared to regions that did not display an increase in CREB1 binding (Figure 7B, middle panel, 7C, Wilcoxon test comparing CRTC1 read counts: p < 2.2 e^-16^), indicating that activity drives both CRTC1 and CREB1 to activity-dependent enhancers.

**Figure 7:**
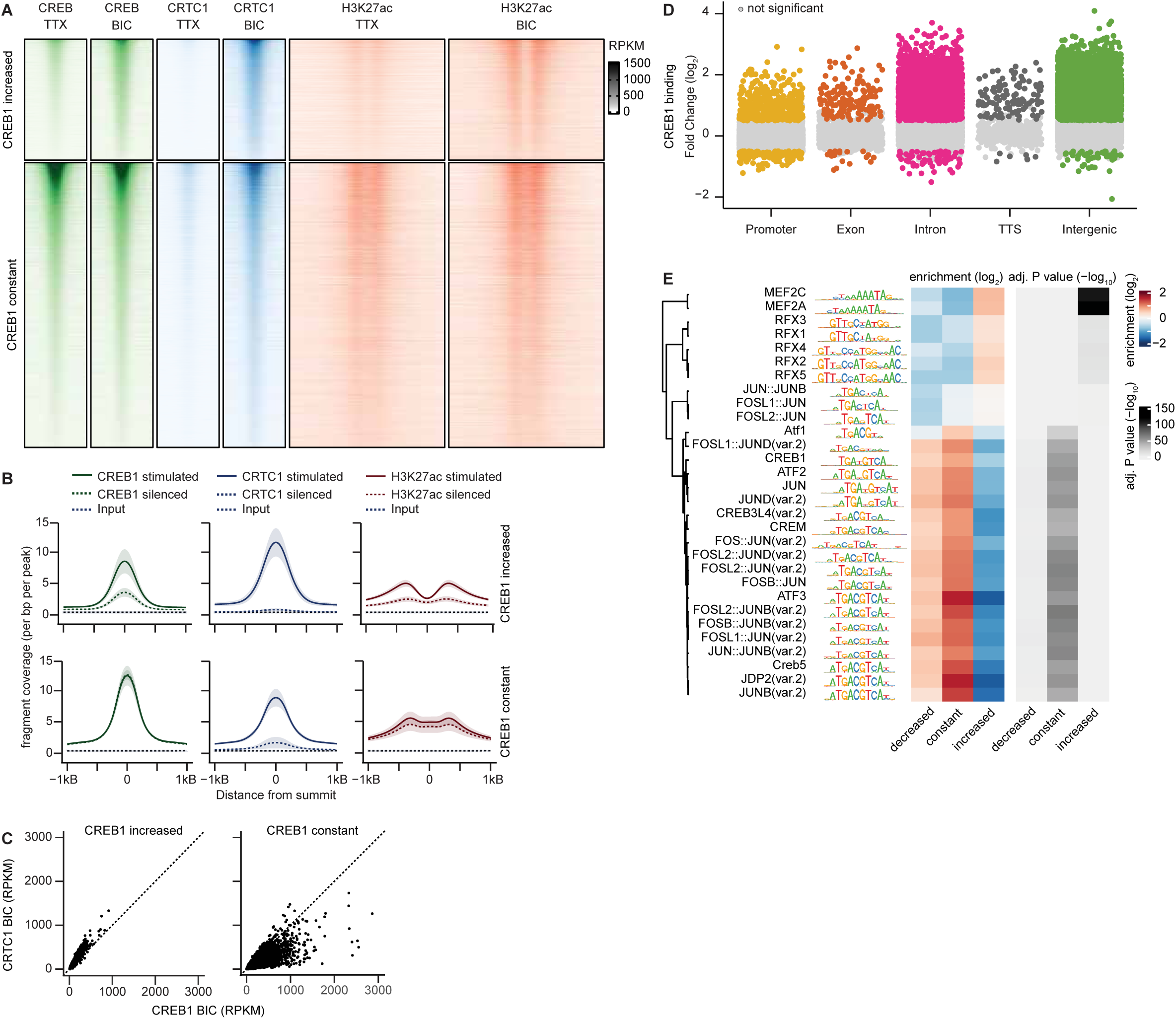
Activity-dependent binding of CRTC1/CREB1 in activity-dependent enhancers containing MEF2C and RFX3 motifs. A) Heatmaps of average normalized ChIP-seq signals for CREB1, CRTC1 and H3K27ac in stimulated and silenced neuronal cultures. Normalized ChIP-seq signals are plotted between +/- 1kB from the peak center for CRTC1 and CREB1 and +/- 2.5 kb for H3K27ac. RPKM = reads per kilobase per million mapped reads. B) ChIP-seq profiles of CREB1, CRTC1 and H3K27ac in regions that show an increase in CREB1 affinity with BIC treatment (upper panel) or regions with constant CREB1 binding with BIC treatment (lower panel). Solid and dashed lines represent average, normalized ChIP-seq signal and shading represents standard deviation. C) Scatter plot of average, normalized ChIP-seq signal of CREB1 and CRTC1 in regions that show an increase in CREB1 affinity with BIC treatment (left) or that show constant CREB1 binding with BIC treatment (right). D) Jitter plot of differential CREB1 binding plotted by genomic annotation. Grey dots represent regions that do not show significant changes in CREB1 affinity with BIC treatment, and colored dots represent regions that show an increase or decrease in CREB1 affinity with BIC treatment. E) Motif enrichment for the 30 most significant motifs identified in CREB1 binding regions that show decreased binding, no change or increased CREB1 binding with BIC treatment.

Consistent with an increase of CREB1 binding at activity-dependent enhancers, the largest increases in CREB1 binding affinity occurred in genomic regions annotated to intronic or intergenic regions, whereas CREB1 binding at promoters remained largely constant (Figure 7D). Motif analysis of regions that either gained, lost or showed constant CREB1 binding revealed that regions with increased CREB1 binding affinity were enriched for MEF2 and RFX motifs, whereas regions with constant CREB1 binding were enriched in motifs for bZIP transcription factors (Figure 7E). Together, these results indicate that in silenced neurons, CREB1 is prebound to promoter and enhancer regions, especially those containing bZIP factor motifs, and that with neuronal activity, CRTC1 and CREB1 are recruited to enhancers that show increased H3K27ac and are enriched in MEF2 and RFX motifs.

### Activity-dependent proximity between CRTC1/CREB1 and MEF2C and RFX3

To further assess the interactions between CRTC1/CREB1 and MEF2C and RFX3, we performed proximity ligation assays (PLA) to visualize interactions between endogenous CRTC1 or CREB1 and endogenous MEF2C and RFX3. With this technique, fluorescent signal is only detected if two candidate proteins are localized within 40 nm of each other^81^. Neurons were either stimulated with 40 µM BIC for 15 min or silenced with 1 µM TTX for 30 min. As a positive control, we first assayed the proximity between CRTC1 and CREB1 and detected PLA signal in stimulated but not in silenced neurons, consistent with the cytoplasmic localization of CRTC1 in TTX-silenced conditions and nuclear localization following BIC-stimulation (Figure 8A, left panel). Next, we tested for interactions between CREB1 or CRTC1 and MEF2C and RFX3. These assays confirmed that both CREB1 (Figure 8A), and CRTC1 (Figure 8B), were in close proximity to both transcription factors following BIC-stimulation. PLA speck count analysis showed that nuclei of stimulated neurons contained a greater number of both CRTC1 and CREB1 interactions with MEF2C and RFX3 compared to silenced neurons. Together, these data reveal activity-dependent contact between CRTC1/CREB1 and MEF2C and RFX3 transcription factors.

**Figure 8:**
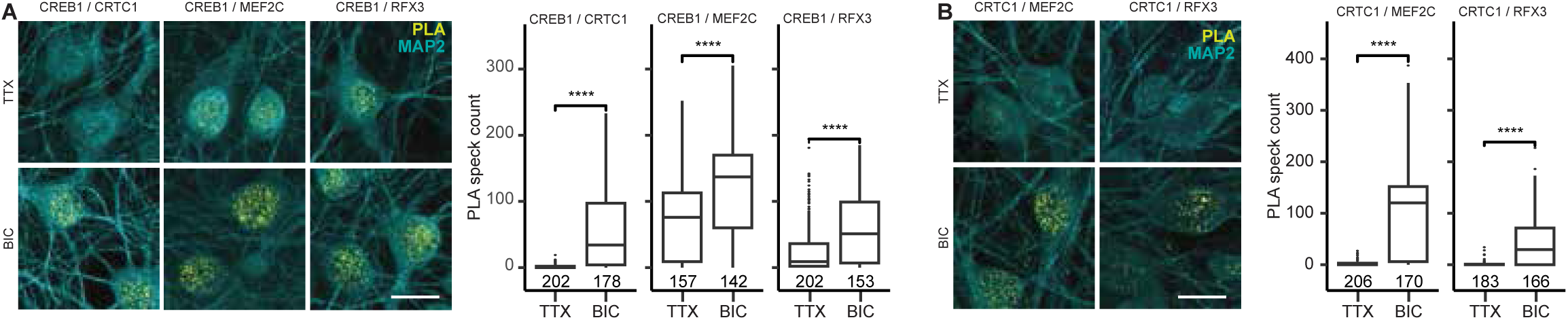
Activity-dependent proximity of CRTC1 and CREB1 to MEF2C and RFX3 in neurons. A) Representative images of PLA of CREB1 with CRTC1, MEF2C or RFX3 in silenced (TTX) and stimulated (BIC) neurons (left). Quantification of PLA signal (right). B) Images of PLA of CRTC1 with MEF2C or RFX3 in silenced (TTX) or stimulated (BIC) neurons (left). Quantification of PLA signal (right). Data were obtained from 3-4 different neuronal cultures. Means of data were compared with a pairwise Wilcoxon test (ns = not significant; p< 0.5 = **; p<0.1 = ***; p<0.01 = ****). Numbers in graphs indicate the number of analyzed cells. Scale bar represents 10 µm.

In summary, our data suggest that synaptic activity facilitates CRTC1 and CREB1 binding to active chromatin and induces combinatorial action between CRTC1/CREB1 and MEF2C or RFX3 at these enhancers. These findings imply that CRTC1/CREB1 coordinate with other factors to regulate activity-dependent gene expression in neurons to support important synaptic functions that are required for the expression of synaptic plasticity.

## Discussion

Glutamatergic stimulation induces gene expression through the activation of various signaling cascades that signal to transcription factors and their cofactors. CREB1 is essential for neuronal survival and long-lasting synaptic plasticity^21^ and is one of the best understood targets of activity-dependent signaling cascades in neurons. Nevertheless, whether and how CREB1 coordinates with other transcription factors is less well explored. Activation of CREB1 regulated transcription occurs in two different ways, by phosphorylation at Ser-133^82,83^ or by recruitment of CRTCs. Phosphorylation of CREB1 occurs through PKA^83,84^, CAMKs^85,86^, MAPK signaling^87,88^ and PI3K/AKT signaling^89,90^. Ser-133 phosphorylation triggers CBP binding, which facilitates transcription^10,11^. CRTCs, on the otherhand, are dephosphorylated by calcineurin and then undergo active import to the nucleus, where they bind CREB1 to activate CREB-dependent transciption^6,8,13,18,56,91^.

Here we used genome-wide CREB1 and CRTC1 profiling in mature rodent forebrain neurons. Consistent with previous reports^12,72–74^, we found that CREB1 was bound to promoters and enhancers in TTX-silenced and BIC-stimulated neurons. CRTC1, in contrast, was recruited to sites of CREB1 binding only in stimulated neurons, consistent with its activity-dependent nuclear translocation. While we identified strongest CREB1 binding activity in promoter regions containing bZIP factor motifs, the majority of CRTC1/CREB1 sites were found in enhancers, albeit at average lower CREB1 occupancy. Further analysis identified activity-dependent increase in CREB1 binding affinity primarily in activity-dependent enhancers that were depleted for bZIP motifs. Interestingly, regions with increased CREB1 occupancy displayed increased CRTC1 recruitment compared to regions that had constant CREB1 binding, suggesting that CRTC1 may support activity-dependent CREB1 binding to these sites. This is consistent with a previous report showing that recruitment of CRTC2 increased CREB1 promoter occupancy in promoters of CREB1 target genes in hepatocytes^92^. Another possibility is that CRTC1 and CREB1 coordinate with other transcription factors, resulting in cooperative binding.

### CRTC1/CREB1 interact with with MEF2C and RFX3

In addition to genome-wide profiling, we employed protein network analysis for CRTC1 using proximity assays and found CRTC1 in proximity to transcription factors other than CREB1, including MEF2 and RFX family members. We also demonstrated that CRTC1 and CREB1 bind to chromatin regions that contain MEF2 and RFX motifs at activity-regulated enhancers. Interestingly, these regions frequently did not contain binding motifs for CREB1. This, however, may not be unexpected, as previous work analyzing genome-wide transcription factor binding profiles demonstrated that most binding events occur in regions that do not contain a matched motif^93^. These binding sites may represent chromatin regions with low affinity or suboptimal binding sequences that require cooperative binding of multiple transcription factors and cofactors for stable chromatin interactions^93^. Indeed, we found that activity-dependent enhancers enriched for MEF2 and RFX motifs have low affinity CREB1 binding in TTX-silenced neurons and BIC-stimulation leads to an increase in CREB1 binding affinity. CRTC1 is a largely disordered protein and intrinsically disordered domains in transcription factors and cofactors have been proposed to facilitate the formation of protein condensates that are involved in transcriptional activation^94^. In agreement with these findings, we found that glutamatergic activity resulted in increased proximity labeling between CRTC1/CREB1 and MEF2C or RFX3. Importantly, CREB1 binding regions with MEF2C motifs were mostly distinct from CREB1 binding regions with RFX3 motifs, indicating that they may regulate different genes.

Previous work has provided evidence that CREB1 and MEF2C co-regulate activity-dependent gene expression. Studies have shown that MEF2C, CREB1 and SRF regulate the activity-dependent expression of *Arc*^95^ and that motifs for these factors are enriched in the promoters of plasticity-related genes^96^. Similarly, neuronal stimulation with Reelin, an extracellular protein that modulates synaptic plasticity, has been shown to trigger an activity-dependent increase in CREB1 and MEF2C binding affinity at Reelin-regulated enhancers^58^.

Recently, RFX family transcription factors have been implicated in activity-dependent gene expression. First, it was reported that enrichment for RFX motifs is greater at activity-dependent enhancers than at promoters^97^. This study also found that a dominant negative mutant of RFX1 primarily binds to enhancers before and after neuronal stimulation^97^. Mutations in RFX family members, including in *Rfx3*, have been found to be associated with autism spectrum disorder (ASD), attention-deficit/hyperactivity disorder and intellectual disability^59^. Additionally, RFX motifs were identified in enhancer regions of ASD risk genes and of genes upregulated in ASD brains. Subsequent work in human iPSC-derived neurons found RFX3 binding at regions upstream of activity-dependent genes and found that RFX3 knockdown resulted in a reduction in activity-dependent gene expression, including in CREB1 target genes. Consistent with our findings, the presence of RFX3 promoted CREB1 binding, suggesting combinatorial action between RFX3 and CREB1 transcriptional action^62^. Likewise, earlier reports demonstrated the involvement of CREB1 and RFX factors in the regulation of MHCII genes in nonneuronal cells^60,61^.

### CRTC1 and mRNA processing in the nucleus

The expression of neuron-specific transcript variants is a hallmark of neuronal differentiation and synaptic plasticity. This is achieved through alternative splicing, alternative polyadenylation or mRNA editing mediated in part by RBPs that are enriched or exclusively expressed in neurons and are regulated by synaptic activity^63,98^. Earlier studies reported that CRTC1 interacts with the splicing factor NONO (p54nrb) to regulate alternative splice site selection of CRTC1/CREB target genes^34,66^. Likewise, overexpression of a constitutively active mutant of CRTC2 led to changes in alternative splicing of about ∼600 transcripts during B-cell differentiation^99^. These data support our APEX2 proximity labeling results in which we identified a significant number of splicing factors and RNA binding proteins within the CRTC1 proxisome. A growing body of recent literature has uncovered the direct involvement of promoters and enhancers and their associated factors in alternative splice site selection and polyadenylation^100^. One such study isolated nuclear proteins bound to newly synthesized RNA in human K652 cells and reported that nearly half of all transcription factors, including CREB1, were bound to RNA^101^. CRTCs were not identified in these studies, likely because they would not be present in the nucleus under these conditions and may not bind RNA directly. Other studies have identified the presence of epigenetic signatures at promoters that denote splice and polyadenylation site selection^102^, so called “dominant promoters”. Promoter dominance was highly prevalent in the *Drosophila* head and in human cerebral organoids, occurring at 40-60% of genes. These epigenetic signatures include enrichment of the CREB1 binding partner CBP/p300 at dominant promoters and their respective 3’-ends, with deletion of CBP leading to a broad dysregulation of 3’-end processing in *Drosophila* embryos. Interestingly, CRTC1 promotes CBP binding to CREB1^103^ and our finding of terminal CRTC1/CREB1 binding sites in some genes raises the possibility of a role for CRTC1/CREB1 in 3’-end selection. Aside from promoter regions, enhancers also modulate 3’-end choice^104^, possibly through the recruitment of RBPs to regions of enhancer-promoter contacts^100^.

### Cytoplasmic CRTC1 and Rho signaling to the cytoskeleton

APEX2 labeling in TTX-silenced neurons revealed proximity between cytoplasmic CRTC1 and various signaling pathways that are important for synaptic plasticity. We identified an extensive set of proteins related to Rho-GTPase signaling and the actin cytoskeleton. Rho-GTPases are regulated by synaptic activity through their GAPs and GEFs and signal to the actin cytoskeleton. Consequently, the outcomes of Rho-GTPase signaling affect all aspects of synaptic spine morphology and dynamics^105^. Interestingly, Rho-GTPases are partially activated through pathways downstream of mTOR. While little is known about the relationship between CRTC1 and Rho-GTPase signaling or the actin cytoskeleton, one such association has been found between p21 activated kinase 4 (PAK4) and CRTC1 in nigral dopaminergic neurons^49^. PAKs are kinases downstream of RAC1/CDC42 signaling and while they are potent regulators of the synaptic cytoskeleton, PAK4 has also been shown to activate CREB1-mediated transcription^106^. Expression of constitutively active PAK4 was found to be neuroprotective in rodent models of Parkinson disease^49^. This study found that PAK4 is in a complex with CRTC1 and CREB1, and that phosphorylation of CRTC1-Ser215 increases the expression of CREB1 target genes and prevents neurodegeneration. Earlier studies have also shown that PAK3 associates in a complex with CRTC1 and CREB1 to facilitate transcription of human T-cell leukemia virus type 1 long terminal repeats^107^. This function of PAKs is unrelated to the direct regulation of the cytoskeleton. Surprisingly, in our APEX2 labeling study, we only identified the association of CRTC1 with PAKs in silenced neurons and not in stimulated neurons, indicating that the CRTC1 – PAK association may function independently from CRTC1’s transcriptional activity.

### CRTC1 and Ras/MAPK signaling

Glutamatergic stimulation activates the Ras/MAPK pathway, which is upstream of activity-dependent gene expression and memory formation^108^. The MAPK signaling pathways includes three kinases, ERK1/2, JNK and p38. In our APEX2 labeling experiment, we mainly identified constituents upstream and downstream of ERK1/2. Phosphorylated ERK1/2 translocates to the nucleus and signals to nuclear targets regulating gene expression, miRNA synthesis and chromatin remodeling^109^. For example, CREB1 is phosphorylated by RSK2 (Rps6ka3) downstream of ERK1/2^87^. Activated ERK1/2 also functions in the cytoplpasm, for example during local translation initiation^110,111^. Direct phosphorylation of CRTC1 through ERK1/2 signaling has not yet been observed, however isoforms CRTC2 and CRTC3 have been shown to be phosphorylated by ERK1/2 directly^112,113^, demonstrating complex regulation of CRTCs and CREB1 by Ras/MAPK signaling.

### CRTC1 and mTOR signaling

Another central regulator of the cellular pathways that underly synaptic plasticity is the protein kinase mTOR^114–116^. mTOR plays a central role during synaptic plasticity through several cellular pathways such as membrane trafficking, translation, cytoskeletal rearrangements and autophagy^117^. As a result, hyperactivity of mTOR is the underlying feature of many neurological diseases^117,118^. mTOR associates with two distinct functional complexes, mTORC1 and mTORC2. mTORC1 regulates protein synthesis^119^ and autophagy^120,121^. mTORC2 on the other hand has been implicated in the regulation of the actin cytoskeleton^122^ and is essential for dendritic spine formation and growth in neurons^123,124^. Within the cytoplasmic CRTC1 proxisome, we identified mTOR and several, but not all, components of mTOR complexes, as well as components of downstream pathways. While it is not known if mTOR phosphorylates CRTC1, it has been shown in the liver that in response to insulin, mTOR phosphorylation of CRTC2 at Ser-136 mediates ER-to-Golgi transport of the transcription factor SREBP1 to regulate lipogenesis^35^. Together, these data and our APEX2 labeling results implicate a general role for CRTCs as mediators of mTOR signaling and the downstream regulation of the secretory pathway, as discussed in more detail in the next section.

### CRTC1 and membrane trafficking

We identified several proteins involved in membrane trafficking in the cyotplasmic CRTC1 proxisome, particularly within the secretory pathway. The secretory pathway is specialized in neurons and is essential for the insertion of membranes and membrane proteins at synapses, and the establishment and maintenance of dendritic structures^125–127^. Previous work has shown that mutations in the secretory pathway result in defects in dendritic complexity in *Drosophila* and rodent neurons^126–128^. As noted earlier, in the liver, CRTC2 regulates ER-to-Golgi transport of SREBP1. This pathway involves CRTC2 directly binding Sec31A, a component of COPII that mediates ER-to-Golgi transport^35^, and this interaction has also been observed in *C. elegans*, which expresses a single CRTC isoform ^31^. After phosphorylation of CRTC2 by mTOR, Sec31A is released and binds to Sec23A, another COPII subunit, to allow SREBP1 transport to the Golgi. We found that cytoplasmic CRTC1 was proximal to Sec31A as well as other COPII subunits and other proteins involved in post-Golgi transport, suggesting that CRTC1 may be associated with the secretory pathway at synapses.

### CRTC1 and translation

Long lasting forms of synaptic plasticity require not only transcription but also translation for their persistence^129–131^. Many transcripts encoding cytosolic and membrane proteins have been shown to localize to synapses^132–135^ and local translation has been demonstrated using fluorescent reporters and metabolic labeling approaches^129,136–140^. To date, the role of CRTC1 in the regulation of translation has not been explored. Within the cytoplasmic CRTC1 proxisome, we identified a large set of translation-related proteins including translation initiation factor EIF2/3/4/5 family members. We also found that CRTC1 is in close proximity to RBPs that regulate localized translation such as CPEBs^51^, FMRP and its interacting proteins CYFIP^52,53^ and PUMs^54^, as well as major signaling pathways that tightly control translation initiation such as mTOR and MAPK signaling networks^50^. Furthermore, translation and processing of membrane proteins requires ER and membrane trafficking, cellular pathways that we also found to be associated with cytoplasmic CRTC1. These findings encourage further investigation of a role for CRTC1 in translational regulation of synaptic membrane proteins.

In excitatory neurons, glutamatergic stimulation leads to the rapid translocation of CRTC1 from the synapse to the nucleus. We show that in neurons, cytoplasmic CRTC1 is part of a larger network of proteins involved in MAPK and Rho signaling, translation and membrane trafficking. After stimulation, we show that CRTC1 translocates to the nucleus where it is in close proximity to a host of transcription factors, chromatin modifiers and splicing machinery that are known to be important for activity-dependent gene expression. We further demonstrate that CREB1 is largely prebound to bZIP motif-containing chromatin regions in neurons, and that CREB1 is selectively recruited to a set of activity-dependent enhancers that are characterized by their preferential CRTC1 recruitment and presence of MEF2 or RFX motifs. Together with our data showing activity-dependent CRTC1/CREB1 association with MEF2C and RFX3, these results point to a model in which cooperative binding of CREB1 with CRTC1, MEF2C and RFX3 at enhancers regulate activity-dependent gene expression in neurons.

## Methods

### Cultured primary neurons

All experimental procedures were approved by the University of California Los Angeles, Animal Research Committee. Forebrains from postnatal day 0-1 Sprague-Dawley rats were dissected in cold HBSS supplemented with 10 mM HEPES buffer and 1 mM sodium pyruvate. Sex was not determined, and tissues from male and female pups were pooled. Our cell culturing protocol results in co-cultures of neurons and glia. The inclusion of glia promotes neuronal health and the formation of mature synapses. In the brain and in our neuronal culture system, CRTC1 expression is restricted to neurons^6,141^. The tissue was sliced and digested with 0.25% Trypsin in HBSS in the presence of 120 mg/mL DNase and 1.2 mM CaCl_2_ for 15 min at 37°C with intermittent mixing. Trypsinization was terminated by washing the tissue twice with DMEM supplemented with 10% fetal bovine serum (FBS). Tissue from one forebrain was resuspended in 2 mL DMEM supplemented with 10% FBS and triturated by passing the solution twelve times through a 1 mL pipet tip. Neuronal cultures were plated at a density of one forebrain for one 10-cm dish in DMEM supplemented with 10% FBS, and medium was exchanged to 25 mL culture medium consisting of Neurobasal A medium supplemented with 1X B-27, 0.5 mM GlutaMax, 25 µM monosodium glutamate, and 25 µM β-mercaptoethanol after one hour. For biochemical assays, cells from one forebrain were plated onto one 10-cm dish containing 1-2 poly-D-lysine-coated coverslips for subsequent fluorescent staining. Cultures were maintained at 37°C and 5% CO_2_ and harvested for experiments at DIV 22.

### Acute hippocampal slices

400-µm thick hippocampal slices were prepared from C57/BL6 mice according to Gray & O’Dell 2013^142^ . In short, hippocampi were dissected in calcium free ACSF (NaCl 124 mM, NaHCO3 25 mM, KCl 4.4 mM, NaH2PO4 1.0 mM, MgSO4 7H2O 1.2 mM, Glucose 10 mM). Per treatment, 5 slices were perfused in interface-type tissue slice chambers at 1 mL/min with warmed, oxygenated (30°C; 95% O_2_ / 5% CO_2_) ACSF containing 2 mM CaCl_2_ for two hours before treatment. Then slices were treated with either normal ACSF or 100 µM DHPG for 10 min by submerging in solution and then immediately flash-frozen in pre-cooled tubes on dry ice and stored at -80°C before processing samples for ChIP-seq.

### Cloning of pLSyn-CRTC1-APEX2

CRTC1-APEX2-FLAG was generated using gene synthesis (Genewiz) and subcloned into pLenti.Syn(0.5).-H2B-eGFP.W (Addgene #51004), replacing H2B-eGFP using the restriction enzymes BstBI and XbaI.

### Lentivirus production and transduction

HEK293T cells were plated into eight T175 flasks at 1x10⁷cells per flask and transfected the next day with pLSyn-CRTC1-APEX2 (6.3 µg DNA), and plasmids encoding Rev (5.55 µg DNA), Gag/Pol (5.55 µg DNA), and VSV-G (1.2 µg DNA) using Polyethylene (PEI) at DNA:PEI ratio of 1:3 to 1:5. The next day, the culture medium was replaced with 30 mL of prewarmed DMEM supplemented with 10% FBS, 10 mM HEPES, 1X Glutamax, and 1 mM sodium pyruvate. After 24 hours, viral supernatant was collected and filtered through a 0.45-µm PES membrane. Virus was pelleted by centrifuging for 2 hours at 22,000 rpm at 4°C and resuspended in 240 µL PBS. The concentrated virus was aliquoted and stored at -80°C. Neurons were transduced at DIV 15 by adding 10 µL of concentrated virus to 5 mL of conditioned medium. After 16 hours, the viral medium was replaced with 20 mL of conditioned medium.

### APEX2 proximity labeling

Samples for stimulated and silenced neurons were prepared separately. Three biological replicates (sets of cultures) were prepared with samples for CRTC1-APEX2 and untransduced controls. For each sample, three 10-cm dishes were used. Neurons were either stimulated with 40 µM BIC and 10 ng/mL Leptomycin B for three hours at 37 °C or silenced with 1 µM TTX for one hour. Previous work demonstrated that 15 min stimulation with BIC leads to a robust nuclear accumulation of endogenously expressed CRTC1^6^, We found that without LMB, CRTC1-APEX2 did not accumulate in the nucleus. Recently, a nuclear export signal was identified in APEX2 that results in biased cytoplasmic localization^143^, which could have been responsible for the inefficient nuclear translocation of CRTC1-APEX2 in the absence of LMB. To label proximal proteins, biotin-phenol was added at a final concentration of 500 µM during the final 30 min of the treatment and labeling was performed by adding H_2_O_2_ at a final concentration of 1 mM for 1 min. Labeling was terminated by washing the neurons three times with quencher solution (PBS with 10 mM sodium azide, 10 mM sodium ascorbate, and 5 mM Trolox (6-hydroxy-2,5,7,8-tetramethylchroman-2-carboxylic acid), supplemented with either 40 µM BIC and 10 ng/mL Leptomycin B, or 1 µM TTX). Cells were lysed for 20 min on ice with RIPA buffer (50 mM Tris pH 7.5, 150 mM NaCl, 0.1% SDS, 0.5% sodium deoxycholate, and 1 % Triton X-100) containing EDTA-free protease inhibitor cocktail (Sigma) and phosphatase inhibitor (PhosSTOP, Sigma), and quenchers (10 mM sodium azide, 10 mM sodium ascorbate, and 5 mM Trolox). Lysates were clarified by centrifugation at 15,000 xg for 10 min at 4°C. Total protein concentration was measured using Pierce 600-nm protein assay reagent (Thermo Fisher Scientific). Samples were concentrated with Amicon centrifugal filter tubes to a protein concentration of >1.5 mg/mL. For each sample, 3 mg of lysate was incubated with 330 µL Pierce streptavidin magnetic beads (Thermo Fisher Scientific) at 4°C overnight. Samples were washed twice with RIPA buffer, once with 1 M KCl, once with 0.1 M Na_2_CO_3_, once with 2 M Urea in 10 mM Tris pH 8, twice with RIPA, and three times with 8 M Urea in 100 mM Tris pH 8.5.

### Liquid Chromatography-Mass Spectrometry sample preparation

Streptavidin beads bound with biotinylated proteins were washed three times each with 8 M urea in 100 mM Tris-HCl (pH 8.5) followed by a wash with nanopure water. Subsequently, beads were resuspended in 100 µL of 50 mM tetraethylammonium bromide. Reduction and alkylation of proteins were performed by sequentially incubating samples with 5 mM tris(2-carboxyethyl)phosphine followed by 10 mM iodoacetamide for 30 min each at room temperature, protected from light, with agitation at 1,000 rpm. Proteins were digested overnight at 37°C with Lys-C (0.4 µg) and trypsin (0.8 µg) under continuous shaking at 1,000 rpm. After digestion, streptavidin beads were removed from the peptide solutions. Peptides were desalted using Pierce C18 tips (100 µL bed volume), dried, and reconstituted in 0.1 % formic acid prior to mass spectrometry analysis.

### Liquid Chromatography-Mass Spectrometry data acquisition

Peptides were separated using a homemade C18 analytical column (75 μm × 25 cm) coupled to a nano-flow Dionex Ultimate 3000 UHPLC system. Peptides were separated using a 140-min gradient of acetonitrile (ACN) at a constant flow rate of 200 nL/min, applied as follows: 1–5.5% ACN from 0– 5 min; linear increase from 5.5–27.5% ACN over 5–128 min; 27.5–35% ACN from 128–135 min; rapid increase to 80% ACN from 135–136 min, maintained at 80% ACN from 136–138 min; followed by a return to 1 % ACN from 138–140 min.

Mass spectrometry analysis was performed using an Orbitrap Fusion Lumos Tribrid instrument operating in data-dependent acquisition mode. Full mass spectrometry spectra were recorded at 120,000 resolution (at 200 m/z), with an automatic gain control target of 2e5 and maximum injection time of 100 ms. MS/MS spectra were acquired at 15,000 resolution after precursor isolation using a window of 1.6 m/z, and fragmentation was performed via higher-energy collisional dissociation at 35% normalized collision energy. A 3-second cycle time was utilized for precursor selection, and dynamic exclusion was set at 25 seconds to prevent repetitive sampling of abundant ions.

### Western blotting

Neurons were lysed in RIPA buffer (50 mM Tris pH 7.5, 150 mM NaCl, 0.1% SDS, 0.5% sodium deoxycholate, and 1% Triton X-100) containing protease and phosphatase inhibitors. Lysates were cleared by centrifugation at 15,000 xg for 10 min and protein concentration was determined by bicinchonic acid protein assay. Lysates were boiled in loading buffer (10% glycerol, 1% SDS, 60 mM Tris-HCl pH 7, 0.1 M DTT, and 0.02% bromophenol blue) for 5 min at 95°C and proteins were separated on 8% polyacrylamide gels. Samples were wet transferred onto a 0.2-µm nitrocellulose membrane and blocked membranes were incubated with mouse anti-FLAG antibody (Millipore-Sigma, F1804) at a dilution of 1:500, followed by incubation with IRDye 800CW-conjugated streptavidin at a dilution of 1:1,000, and anti-mouse IRDye 680CW antibody at a dilution of 1:10,000. Membranes were imaged using the Odyssey infrared imaging system (LI-COR Biosciences).

### Fluorescence staining and imaging of neuronal cultures

Coverslips from 10-cm dishes that were used for biochemical experiments were transferred to 24-well dishes using forceps. Neurons were fixed for 10 min with 4% paraformaldehyde at room temperature and solubilized with 0.1% Triton X-100 for 5 min. Neurons were stained with rabbit anti-CRTC1 antibody (Bethyl, A300-769A) at a dilution of 1:1,00 and mouse anti-FLAG antibody (Millipore-Sigma, F1804-200UG) at a dilution of 1:500, followed by secondary antibodies goat anti-rabbit Alexa 647, goat anti-mouse Alexa 488 and streptavidin-Alexa 555 at a dilution of 1:1,00. Confocal fluorescent images were taken on a Zeiss LSM880 system.

### Fluorescence staining and imaging of hippocampal slices

Immediately after treatment, slices were fixed overnight in 4% paraformaldehyde. Slices were washed three times with PBS, covered in HistoGel and embedded in paraffin. 4-µm thick paraffin sections were serially rehydrated and deparaffinized. After heat-induced antigen retrieval, cell membranes were permeabilized with 0.1% Triton-X-100 at room temperature for 30 min, then blocked in 10 % goat serum at room temperature for one hour. Slices were then incubated in primary antibodies (anti-CRTC1 (Bethyl A300-769A,1:500), anti-MAP2 (Phospho Solutions1100-MAP2, 1:1,00) overnight at 4°C and with secondary antibodies and Hoechst 33342 (Thermo Fisher Scientific H3570) for 2 hours at room temperature. Confocal fluorescent images were taken on a Zeiss LSM700 system.

### Proximity Ligation Assays

Neurons were silenced with 1 µM TTX for 30 min or stimulated with 40 µM BIC for 15 min at 37°C and fixed with 4 % paraformaldehyde in PBS for 10 min. Neurons were permeabilized with 0.25 % Triton X-100 for 15 min, washed 3 times with PBS, and blocked with Duolink blocking buffer for 1 hour at 37°C. Neurons were then incubated with primary antibodies against CRTC1 (Bethyl A300-769A,1:500), mouse CREB1 (Cell signaling 9104S, 1:500), rabbit CREB1(Cell signaling 9197S, 1:1,00) MEF2C (Thermo Fisher Scientific MA5-25477, 1:2000), RFX3 (Thermo Fisher Scientific TA505916, 1:500), and MAP2 (Phospho Solutions1100-MAP2, 1:1,00) overnight at 4°C. Neurons were washed with Duolink washing buffer A twice for 5 min and incubated with Duolink probes PLUS and MINUS for 1 hour at 37°C inside a humidity chamber. After washing with Buffer A twice for 5 min, probes were ligated for 30 min at 37°C. Neurons were washed with buffer A twice for 5 min and incubated with polymerase and Duolink Orange Detection Reagent for 100 min at 37°C. Neurons were washed with wash buffer B supplemented with secondary antibody goat anti-chicken Alexa 647 (1:1,00) and Hoechst dye 33342 (1:1,00) twice for 15 min. Neurons were washed with PBS three times for 5 min and coverslips were mounted with Aqua-Poly/Mount. For quantification of PLA signals, Z-stacks confocal images were taken on a Zeiss LSM880 with a 63X objective. Z-stacks were collapsed with maximum intensity projection using ImageJ and cells and nuclei were delineated based on MAP2 and Hoechst staining, respectively. The number of PLA speckles were counted within the respective compartments using find maxima function of ImageJ with prominence of 25. Speck counts per cell were plotted with the R package ggplot2 and means were compared using the stat_compare_means() function with a pairwise Wilcoxon Test (wilcox.test). For each condition, cells from 3-4 separate cultures were analyzed. Numbers of analyzed cells are indicated in the figure.

### ChromatinIP from cultured neurons and library construction

Neuronal cultures were treated for 15 min with 40 µM BIC or for 30 min with 1 µM TTX. Cultures were washed with Tyrode’s solution (140 mM NaCl, 10 mM HEPES pH 7.4, 5 mM KCl, 3 mM CaCl_2_, 1 mM MgCl_2_, 20 mM glucose) and fixed with 2 mM DSG in Tyrode’s solution for 25 min, followed by 1 % formaldehyde for 10 min. Fixatives were quenched with glycine at a final concentration of 150 mM for 5 min. Cells were collected with a cell scraper and pelleted at 1,00 xg for 5 min and stored at - 80°C until further use. Cell pellets from approximately ten 10-cm dishes were resuspended in 2 mL sonication buffer (50 mM HEPES pH 7.4, 140 mM NaCl, 1 mM EDTA, 1 % Triton X-100, 0.1% sodium deoxycholate, 0.1 % SDS) and chromatin was fragmented using a Misonix S-4000 probe sonicator (1/8 probe tapered 419, 6 min, 3 sec on, 15 sec off at 30% power). Debris was removed by centrifugation and the DNA concentration of the sonicate was determined using the Qubit dsDNA BR Assay Kit. For each condition, two IPs were performed (CRTC1/CREB1: 15 µg chromatin; H3K27ac: 10 µg chromatin, H3K4me3: 5 µg chromatin) and combined during washes. Antibodies were diluted in PBS (anti-CRTC1: 30 µg, anti-CREB1: 10 µ<u>l (concentration unknown)</u>, H3K27ac: 5 µg, H3K4me3: 5 µg) and bound to 25 µL washed magnetic Protein G beads (Active Motif) for 4 hours at 4 °C. Chromatin was precipitated overnight at 4 °C and beads were washed twice with wash buffer I-IV (I: 50 mM Tris-HCl pH 8, 150 mM NaCl, 2 mM EDTA, 1% NP-40, 0.1% sodium deoxycholate, 0.1% SDS; II: 20 mM Tris-HCl pH 8, 150 mM NaCl, 1 % Triton X-100, 0.1 % SDS; III: 20 mM Tris HCl pH 8, 500 mM NaCl, 2 mM EDTA, 1 % Triton X-100, 0.1% SDS; IV: 10 mM Tris-HCl pH 8, 250 mM LiCl, 1 mM EDTA, 1% sodium deoxycholate, 1% NP-40), once with TE buffer and then resuspended in TE buffer. DNA was extracted by incubation with RNase A for 5 min at 37 °C and Proteinase K over night at 60 °C followed by phenol chloroform extraction and ethanol precipitation. Four biological replicates (sets of cultures) were generated for each treatment, and aliquots from the same sonicate were used for CRTC1 and CREB1, and H3K4me3 and H3K27ac. Sequencing libraries were generated from precipitated DNA or 10 ng of fragmented input DNA using the Next Generation Sequencing DNA Library Kit (Active Motif) following the guidelines for > 10 ng input and 200 bp fragments. Libraries were PCR amplified with 12 cycles. Quality and size distribution of libraries were verified by D1000 Screentape (Agilent) and library concentration was determined with Qubit dsDNA HS (high sensitivity) Assays. Libraries were sequenced to a depth of ∼ 20 - 50 million reads per sample.

### Chromatin IP from CA1 and library construction

Hippocampal CA1 was dissected in Tris-buffered saline containing phosphatase and protease inhibitors and submerged immediately in 2 mM DSG in low binding tubes and fixed for 25 min at room temperature (4 slices per tube). Then the solution was replaced with 1 % formaldehyde for further fixation for 10 min. Fixatives were quenched with glycine (final concentration of 125 mM) for 10 min and samples were kept on ice until all samples were processed. Tissue was washed twice with PBS and 20 CA1s were pooled per condition per biological replicate. Tissue was lysed in NP-40 buffer (50 mM HEPES, pH 7.4, 150 mM NaCl, 1% NP-40, 10 mM EDTA, protease and phosphatase inhibitors) for 15 min with intermittent vortexing, followed by sonication in 500 µL sonication buffer (50 mM Tris-HCl, pH 8.0, 1% SDS, 10 mM EDTA, protease and phosphatase inhibitors). Chromatin was fragmented using a Misonix S-3000 probe sonicator (1/8 probe tapered 419, 2:30 min, 3 sec on, 15 sec off, Power 5.5). Debris was removed by centrifugation at maximum speed for 10 min. Chromatin IP was performed with the Low Cell ChIP kit (Active Motif) according to the manufacturer’s instructions with modifications described below. We included unique molecular identifiers (UMIs) in the library preparation to allow more accurate PCR duplicate removal. Sonicate was diluted 10-fold to 5 mL to reduce the concentration of SDS before IP (dilution buffer: 16.7 mM Tris-HCl, pH 8.1, 167 mM NaCl, 1.2 mM EDTA, 0.01 % SDS, protease and phosphatase inhibitors). Each sample was split into 4 aliquots for IP. Per aliquot, 2.5 µg of CRTC1 antibody was used for a total of 10 µg/ per sample. IP was performed overnight at 4 °C. Input was phenol-chloroform purified DNA from sonicate. ChIP DNA was stored at -80 °C until all samples were acquired. Sequencing libraries were generated from precipitated DNA or 10 ng fragmented input DNA using the Next Generation Sequencing DNA Library Kit (Active Motif) following the guidelines for <10 ng input and 200 bp fragments. Libraries were PCR amplified with 12 cycles. Quality and size distribution of libraries were verified by D1000 Screentape (Agilent) and library concentration was determined with Qubit dsDNA HS (high sensitivity) Assays. Three biological replicates (slices from separate mice on separate experimental days) for ChIP-seq were generated for each condition. Libraries were sequenced to a depth of ∼ 40 - 50 million reads per sample.

### Liquid Chromatography-Mass Spectrometry data analysis

MS/MS data were analyzed using MaxQuant software (version 1.6.17.0) by searching against the rat reference proteome from EMBL (UP000002494_10116_RAT_Rattus norvegicus, 22401 entries). Carbamidomethylation of cysteine residues was specified as a fixed modification, whereas oxidation of methionine and N-terminal acetylation were set as variable modifications. Trypsin was selected as the digestion enzyme, allowing up to two missed cleavages. Precursor mass tolerances were set to 20 ppm for the initial search and 4.5 ppm for the subsequent main search, while fragment mass tolerance was set to 20 ppm. Identification results were filtered using a false discovery rate threshold of 1 % at both peptide spectral match (PSM) and protein levels.

Peptide quantification was conducted using MaxQuant’s label-free quantification (LFQ) algorithm. Differential protein enrichment in the CRTC1-APEX2 proximity labeling experiment was assessed statistically using MSstats (version 3.10). LFQ intensities were normalized via equalized medians, and protein-level summarization employed the Tukey median polish method. Statistical significance (p-values from t-tests) was adjusted for multiple hypothesis testing using the Benjamini–Hochberg procedure. Statistically significant proteins were selected as iP-value < 0.05, iLog_2_FC > 2, and present in all three replicates.

### Subcellular localization, Gene Ontology analysis and Protein Interaction networks

Subcellular localization of proteins was determined using the COMPARTMENTS database (https://compartments.jensenlab.org/)^41^ Data were based on known localizations of human proteins. Gene Ontology analysis was performed using Enrichr (https://maayanlab.cloud/Enrichr/)^42–44^. High confidence, known protein interactions of CRTC1 proximal proteins were visualized using Cytoscape and human STRING database with a confidence (score) cutoff of 0.99. Simplified, functional categories were manually assigned using STRING^47^ descriptions and Genecards^45,46^ as guidance.

### ChIP-seq data processing and peak calling for cultured neurons

Sequencing results were demultiplexed and PCR duplicates were removed based on UMIs using custom software (High-Throughput Sequencing Core, Broad Stem Cell Research Center, UCLA) with a hamming distance of 1. Reads were aligned to rn6 using Bowtie2^144^. Blacklisted regions were not determined for the rat genome^145^, therefore we generated a list of high signal regions in input read files. First genome coverage per basepair was calculated using bedtools genomecov^146^. Average coverage was 3. Regions with a count of > 200 were selected and regions within 15,000 bp were merged using bedtools merge. All regions were visually inspected to define region borders using IGV Viewer. Blacklisted regions were removed from read files before further analysis using bedtools intersect. Effective genome size for the rn6 genome was calculated using faCount -summary from Kent tools (UCSC genome browser) and was determined as the number of non-N bases = 2.65e9. We observed several ChIP-seq peaks with multiple summits in ChIP samples for CRTC1 and CREB1, therefore peak summits were called with macs2 callpeak^71,147^ with the options -B --SPMR, -g 2.65e9, --keep-dup all, --call-summits, -q 0.05. For histone marks peaks were called with --broad --broad-cutoff 0.05. Consensus peak sets were generated with DiffBind^148^. For CRTC1 and CREB1, read files from each sample were subsampled to 20 million reads with samttols view –s option and merged per factor and condition using samtools merge. Peak summits were called on merged read files and extended by 100 bp to each direction using bedtools slop. A file of consensus peak summits was obtained by requiring the peak summits to be present in all four replicates within +/- 100 bp of the peak summit called on the merged read files. The position of peak summits obtained from merged read files was defined to be the consensus peak summit. For histone marks, consensus peaks were required to be present in all four replicates.

### ChIP-seq data processing and peak calling for CA1

Sequencing results were demultiplexed and PCR duplicates were removed based on UMIs using custom software (High-Throughput Sequencing Core, Broad Stem Cell Research Center, UCLA) with a hamming distance of 1. Reads were aligned to mm10 using Bowtie2. Blacklisted regions were obtained from ENCODE^145^. More artefact regions of high signal in input samples were identified using GreyListChIP^149^ and manual annotation. Artefact regions were removed using bedtools intersect. Peaks were called using macs2 callpeak with the options -B --SPMR, -g mm, --keep-dup all, --call-summits, -p 0.01. Consensus peak sets were generated using DiffBind with the requirement that peaks had to be present in all replicates per condition.

### Downstream analysis of ChIP-seq data

Genomic annotation and metagene profile plots were determined with HOMER^150^ annotatePeaks.pl. BigWig files were generated with Deeptools^151^ bamCoverage (options --binSize 10 --normalizeUsing RPKM --ignoreForNormalization chrX --extendReads 300, rat: --effectiveGenomeSize 2651731494 or mouse: --effectiveGenomeSize 2913022398. BigWig files were used as input for the Enriched Heatmaps^152^ function normalizeToMatrix() with the option w=10 and averages of all replicates were generated with getSignalsFromList() and plotted with EnrichedHeatmap(). Browser tracks were generated from normalized macs2 bedgraph files using Gviz^153^. Motif enrichment was performed with monaLisa^75^ with the function calcBinnedMotifEnrR() (options: background = "genome", genome = BSgenome.Rnorvegicus.UCSC.rn6, genome.regions = NULL, # sample from full genome, genome.oversample = 2). Motif scan was performed with findMotifHits() (options: min.score = 10, method = "matchPWM"). Position weight matrices (pwms) were obtained from JASPAR with getMatrixSet(JASPAR2020, opts = list(matrixtype = "PWM",tax_group = "vertebrates")). Peak set overlap was calculated with bedtools intersect. Data were processed with R and remaining graphs were plotted with ggplot2^154^.

## Acknowledgements

The authors would like to thank the UCLA Broad Stem Cell Research Center Sequencing Core for data collection and the UCLA Quantitative C Computational Biosciences Collaboratory for data analysis support, and funding from the David Geffen Foundation (to KCM) and NIH R01 MH07722 (to KCM).

## Notes

### Competing Interest Statement

The authors have declared no competing interest.

